# Central and peripheral innervation patterns of defined axial motor units in larval zebrafish

**DOI:** 10.1101/559062

**Authors:** Saul Bello-Rojas, Ana E. Istrate, Sandeep Kishore, David L. McLean

## Abstract

Spinal motor neurons and the peripheral muscle fibers they innervate form discrete motor units that execute movements of varying force and speed. Subsets of spinal motor neurons also exhibit axon collaterals that influence motor output centrally. Here, we have used *in vivo* imaging to anatomically characterize the central and peripheral innervation patterns of axial motor units in larval zebrafish. Using early born ‘primary’ motor neurons and their division of epaxial and hypaxial muscle into four distinct quadrants as a reference, we define three distinct types of later born ‘secondary’ motor units. The largest are ‘m-type’ units, which innervate deeper fast-twitch muscle fibers via medial nerves. Next in size are ‘ms-type’ secondaries, which innervate superficial fast-twitch and slow fibers via medial and septal nerves, followed by ‘s-type’ units, which exclusively innervate superficial slow muscle fibers via septal nerves. All types of secondaries innervate up to four axial quadrants. Central axon collaterals are found in subsets of primaries based on soma position and predominantly in secondary fast-twitch units (m, ms) with increasing likelihood based on number of quadrants innervated. Collaterals are labeled by synaptophysin-tagged fluorescent proteins, but not PSD95, consistent with their output function. Also, PSD95 dendrite labeling reveals that larger motor units receive more excitatory synaptic input. Collaterals are largely restricted to the neuropil, however perisomatic connections are observed between motor units. These observations suggest that recurrent interactions are dominated by motor neurons recruited during stronger movements and set the stage for functional investigations of recurrent motor circuitry in larval zebrafish.

## 1. Introduction

Motor neurons are the so-called ‘final common pathway’ for a range of behaviors in all mobile animals (Sherrington, 1904). In vertebrates, different types of motor neurons and the muscle fibers they innervate, collectively known as motor units, generate movements of different forces and speeds (Denny-Brown, 1929; Eccles et al., 1958). In general, motor units characterized as ‘fast’ are responsible for delivering maximum, but transient speeds and forces, while those characterized as ‘slow’ deliver minimum, but sustained speeds and forces (Fetcho, 1992; Heckman and Enoka, 2012). Faster motor units also tend to be larger in terms of motor neuron and muscle fiber size, and the number of fibers innervated (Burke, 1981; Henneman and Mendell, 1981).

Another common feature of spinal motor neurons in a variety of animals is the presence of axon collaterals that branch off prior to exiting the ventral root (Prestige, 1966; Chmykhova et al., 2005). This means that some motor neurons are poised not only to generate peripheral muscle contractions, but also to generate central commands. Physiological studies have revealed that stimulation of motor neurons can yield both excitatory and inhibitory effects on motor output (Renshaw, 1941; Eccles et al., 1954; Perrins and Roberts, 1995; Mentis et al., 2005; Nishimaru et al., 2005; Bhumbra et al., 2014; Moore et al., 2015; d’Incamps et al., 2017; Bhumbra and Beato, 2018). These effects are most likely explained by recurrent excitation of neighboring motor neurons and inhibitory Renshaw cells (Cullheim et al., 1977; Chmykhova and Babalian, 1993; Adanina et al., 2005). However, despite their ubiquity and history of investigation the function of recurrent motor circuits is still debated (Hultborn et al., 1979; Lindsay and Binder, 1991; Maltenfort et al., 1998; Alvarez and Fyffe, 2007; Obeidat et al., 2014).

To better understand the origins of recurrent motor circuits and their potential function, we have anatomically assessed the central and peripheral innervation patterns of spinal motor neurons in larval zebrafish. Larval zebrafish swim primarily using their axial musculature, which can generate tetanic forces between 200–400 mN/mm^2^ (Martin et al., 2015) and tail-beat frequencies between 20–100 Hz (Muller and van Leeuwen, 2004). To do so, they use fast-twitch, ‘white’ fibers that make up the bulk of the muscle mass and a superficial layer of slow, ‘red’ fibers (Devoto et al., 1996; Coutts et al., 2006; Luna et al., 2015). At faster speeds, axial motor neurons are recruited in a topographic pattern from the bottom of spinal cord up (McLean et al., 2007; Knafo et al., 2017), with smaller, ventral motor neurons innervating slow muscle fibers recruited before larger, dorsal motor neurons innervating fast-twitch muscle (Menelaou and McLean, 2012; Wang and Brehm, 2017).

In cats, where motor units are well-defined based on innervation patterns, there is evidence for interactions both within and between distinct types (Cullheim and Kellerth, 1978; Friedman et al., 1981; Cullheim et al., 1984; Hultborn et al., 1988a; Hultborn et al., 1988b; McCurdy and Hamm, 1994; Turkin et al., 1998). In larval zebrafish, axial motor neurons also exhibit central axon collaterals (Menelaou and McLean, 2012), however the relationship between motor unit identity, the presence or absence of central collaterals, and their potential targets has not been explored. Thus, our goal here was three-fold: 1) to better characterize differences in the innervation patterns of axial motor neurons to help define distinct axial motor units; 2) to link the existence of central axon collaterals to motor unit identity; and, 3) to identify the potential targets of central collaterals.

## 2. Materials and methods

### Wildtype and transgenic fish

Adult zebrafish were maintained at 28.5°C on a 14/10-hour light/dark schedule in a custom-built facility (Aquatic Habitats). Zebrafish embryos were obtained from daily crosses of adults and were raised at 28.5 °C. Experiments were performed using 5-7-day-old larval zebrafish when they have a fully inflated swim bladder and are free-swimming. At this developmental stage, zebrafish have not yet sexually differentiated and are nourished by their yolk. All procedures described below conform to NIH guidelines regarding animal experimentation and were approved by the Northwestern University Institutional Animal Care and Use Committee.

To generate stable transgenic fish expressing green fluorescent (GFP) in motor neurons, we injected a single plasmid into wild type embryos at the single cell stage along with *in vitro* transcribed transposase mRNA. This plasmid contained a Tol2 destination vector (pDest-Tol2pA2; Kawakami et al., 1998) with Tol2 transposable elements flanking three copies of the 125 basepair (bp) mnx1 enhancer (Zelenchuk and Bruses, 2011) followed by prenylated GFP and a SV40 polyadenylation (pA) signal. As a short hand, we will refer to these as Tg[mnx1:GFP] fish. To generate transgenic fish expressing red fluorescent protein (RFP) in motor neurons we co-injecting two plasmids with *in vitro* transcribed transposase mRNA. The first plasmid was a Tol2 destination vector with Tol2 transposable elements flanking three copies of the 125 bp mnx1 enhancer, followed by Gal4-VP16 (Sadowski et al., 1988) and a SV40 pA signal. The second plasmid was a Tol2 destination vector with Tol2 transposable elements flanking five copies of the UAS promoter, followed by pTagRFP (Evrogen), and a SV40 pA signal. We will refer to these as Tg[mnx1:Gal4;UAS:pTagFRP] compound fish.

For injections to create stable lines, the concentration of injected plasmids was 20 ng/μl and the concentration of the transposase mRNA was 50 ng/μl. Messenger RNA was synthesized using the mMessage mMachine kit (Ambion). Following injections, putative founder embryos (F0) were then raised to sexual maturity and crossed with widtype adults to generate transgenic F1 embryos that were screened for red or green fluorescence at 36 hours post fertilization (hpf).

For transient labeling we used mnx:GFP or the Gal-UAS system to drive mosaic expression of reporter constructs selectively in motor neurons (Koster and Fraser, 2001; Seredick et al., 2012). Gal4 was driven by the zebrafish mnx1 gene (mnx1:Gal4) or the bacterial artificial chromosome for vesicle acetylcholine transporter (VAChT:Gal4) (Kishore and Fetcho, 2013). Reporter constructs containing upstream activating sequences (UAS) included membrane-associated fluorescent proteins, mCD8:GFP and pTagRFP (Menelaou et al., 2014), and synapse-specific fluorescent proteins Syp:GFP (Meyer and Smith, 2006), Syp-pTagRFP (Menelaou et al., 2014), and PSD95:GFP (Niell et al., 2004). Mosaic motor neuron expression was obtained by co-injecting either mnx:GFP or a combination of either mnx1:Gal4 or VACht:Gal4 plasmid with different combinations of the reporter constructs into one- to two-cell stage wild-type embryos or Tg[mnx1Gal4;UAS:pTagRFP] embryos using a microinjector (Narishige model IM300). For transient injections, DNA solutions were prepared at concentrations between 10 and 20 ng/*µ*l.

### Confocal imaging and morphological analysis

Zebrafish larvae were anesthetized in 0.02% w/v ethyl 3-amino-benzoate methanesulfonic acid (MS-222; Sigma-Aldrich), placed in a glass-bottomed dish, and embedded on their side in low-melting-point agar (1.4% in system water). Once the agar had solidified, more anesthetic solution was added to prevent agar desiccation and movement of the fish. Images were acquired with a confocal microscope (Zeiss LSM 710) equipped with an argon laser (488 nm) and a Helium/Neon laser (543 nm) using a Zeiss 40x/1.0-NA or 20x/1.0-NA water-immersion objective. For any given motor neuron, one z-stack was acquired to capture the entire morphology of the cell using the 20x objective with an optical section of 2 *µ*m. A second z-stack was acquired to capture at morphology within spinal cord at higher resolution using the 40x objective with an optical section of 0.70 *µ*m. Motor neurons sampled were confined to muscle segments 5-22 or ‘midbody’. In 23 of 303 motor neurons, only lower magnification images were collected so we could not perform statistical analysis of intraspinal features.

Three-dimensional (3D) images were reconstructed and analyzed using Imaris (Bitplane). Axon and dendrite length measurements of each reconstructed motor neuron was obtained using the Filament function to trace over the 3D rendering. Total peripheral innervation measurements included the main axon length within the spinal cord and the total length within the musculature. Central collateral length included only extensions that sprouted from the main axon within the spinal cord. The center of the soma was used as the point of origin for tracing the dendrite filament. Detection of both central and peripheral putative synaptophysin and central PSD95 boutons was performed by using the Spots function and a threshold value of 1 *µ*m. Coordinates were then exported from Imaris and plotted using previously published Matlab scripts (Kishore and Fetcho, 2013) interfaced with Imaris to display the position of individual filament nodes and synaptophysin and PSD95 puncta. To account for variations in spinal cord height and width across different fish, these coordinates were normalized relative to the dorsal and edges of the spinal cord as well as along medio-lateral landmarks marked medially by the ipsilateral Mauthner axon, and laterally by the lateral-most boundary between the spinal cord and the axial musculature. Nomalized coordinates from all motor neurons or within sub-types of motor neurons were then pooled to generate separate contour plots that represent the variation in densities of filament coordinates or synaptophysin or PSD95 puncta. Distribution contours were made in Matlab using the kde2d function (Matlab File Exchange), which estimates a bivariate kernel density over the set of normalized coordinates using a 32 by 32 grid. (Botev et al., 2010; Bikoff et al., 2016). Contour plots were then generated from the calculated densities using the “contour” function in Matlab. Contour lines are displayed in increasing increments of one-sixth of the peak density where the innermost contour represents the highest density of coordinates.

Dorso-ventral soma positions were measured from the center of the cell body and normalized to a measurement of the dorsal and ventral edges of the spinal cord from differential interference contrast (DIC) images captured during image acquisition on the confocal microscope. The peak dorso-ventral height of the dendrites and axon collaterals were measured using the same method. Rostro-caudal position was normalized to the ventral root by setting the exit point as the 0 *µ*m mark. Any position located rostral to the ventral root was positive, while anything caudal to the ventral root was negative. Soma size was quantified by measuring the diameter at its widest point in the DIC image. Medio-lateral positions were normalized to the ipsilateral Mauthner axon and the lateral-most boundary between spinal cord and the axial musculature.

### Statistical analysis

All data were first tested for normality before selecting the appropriate parametric or non-parametric test. Statistical tests, their critical values, degrees of freedom, and significance are reported where appropriate in the main text. Statistical analysis was performed using Microsoft Excel with a StatPlus plug-in (AnalystSoft). All data are reported as means plus/minus standard deviations.

## 3. Results

### Secondary motor neurons can be defined based on the identity and amount of target musculature

Previous studies in larval zebrafish have divided axial motor neurons into two groups. ‘Primary’ motor neurons are the first to develop and innervate non-overlapping regions of dorsal epaxial and ventral hypaxial musculature (Myers et al., 1986; Menelaou and McLean, 2012). ‘Secondary’ motor neurons develop next and can innervate distinct or overlapping regions of epaxial and hypaxial musculature (Myers, 1985; Menelaou and McLean, 2012). While secondary motor neurons have been characterized based on electrophysiological properties and/or main axon trajectories (Menelaou and McLean, 2012; Asakawa et al., 2013), to better resolve the likelihood of targeting peripheral slow *versus* deeper fast-twitch muscle fibers and the extent of that innervation, we turned to *in vivo* confocal microscopy.

To begin exploring the patterns of peripheral innervation, we created two transgenic lines using a promoter for the motor neuron specific transcription factor, mnx1 (Zelenchuk and Bruses, 2011; Seredick et al., 2012), which is a homolog of mammalian Hb9 (Arber et al., 1999; Thaler et al., 1999; Ferrier et al., 2001). For purposes of analysis, we focused on neurons at midbody (segments 5-22) to avoid rostral pectoral fin motor neurons and caudal fin motor neurons (Figure 1a,b). In 5-7 day old Tg[mnx1:GFP] and Tg[mnx1:Gal4;UAS:pTagRFP] larvae, both the somata and peripheral axons of spinal motor neurons can be observed (Figure 1a). From the transverse perspective, peripheral motor nerves arising from motor neuron somata at midbody exit spinal cord and immediately divide into dorsal and ventral branches, from which medial nerves branch within the deeper fast-twitch muscle and septal nerves run superficially to innervate slow muscle (Figure 1c,d), matching the innervation patterns of axial muscles described in adult zebrafish (Westerfield et al., 1986). This pattern of innervation was observed along the body, although the height and width of the trunk decreased in more caudal regions (Figure 1d).

**Figure 1.**
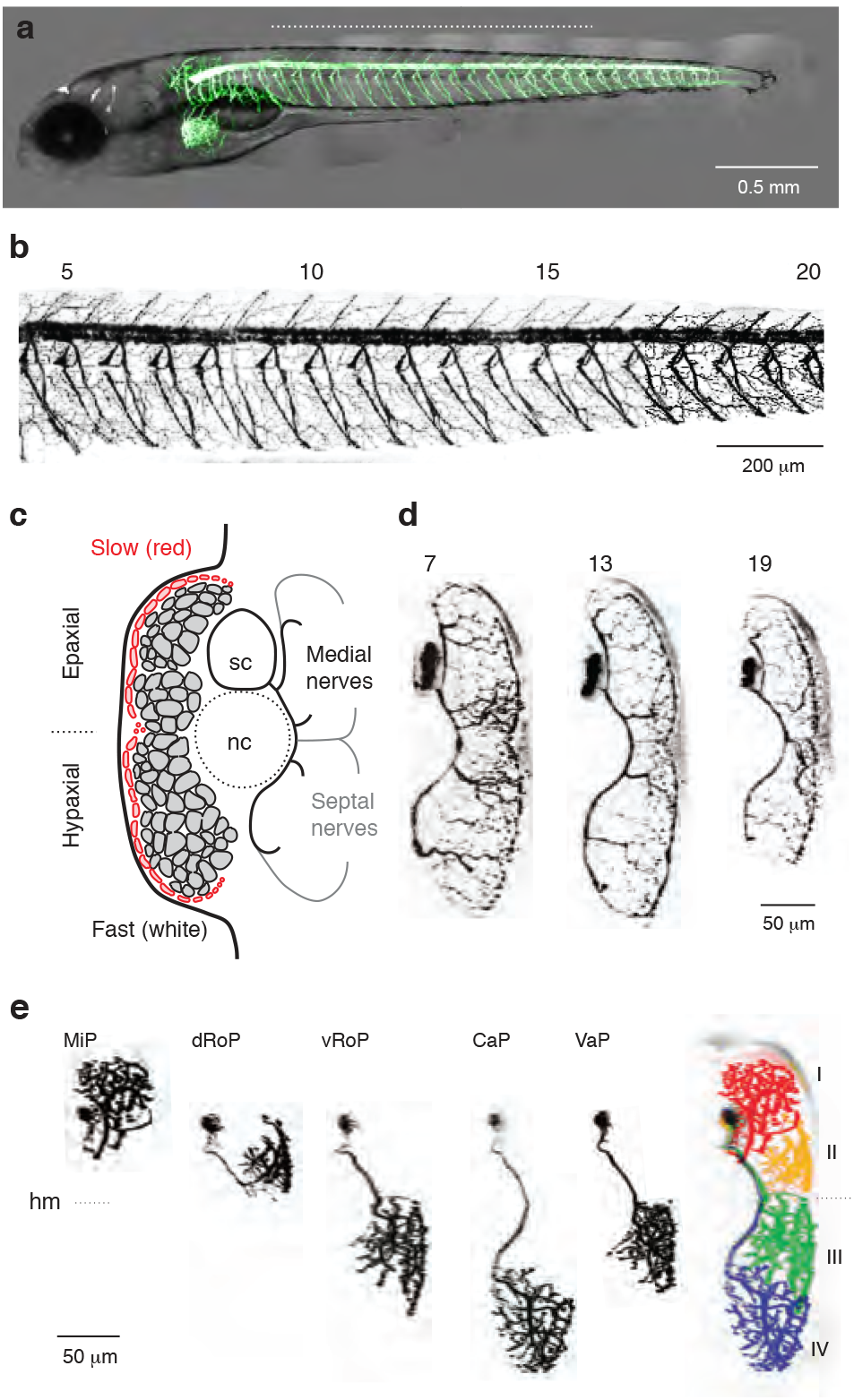
Visualizing the organization of axial muscle innervation in larval zebrafish. **a,** Side view of 5 day old Tg[mnx1:GFP] zebrafish illustrating spinal motor neurons and peripheral motor nerves in green/white. Rostral is to the left and dorsal is up. Dashed white line indicates region expanded in *b*. Note, some fluorescence is also observed in the brain. **b,** Contrast-inverted image of axial motor neurons and peripheral motor nerves between the 5^th^ and 20^th^ body segment (numbered) viewed from the side. Motor neurons form a longitudinal column near the top of the image. **c,** Schematic illustrating the medio-lateral distribution of fast-twitch muscle fibers (grey) and slow muscle fibers (red) in transverse view. Dorsal epaxial and ventral hypaxial musculature is separated by the horizontal myoseptum (black dashed horizonal line). Peripheral nerves originating from the spinal cord (sc) are distinguished by their medial (black) and septal (grey) projection patterns. Dashed circle indicates the notochord (nc). **d**, Unilateral transverse views taken from different body segments (numbered). **e**, Contrast-inverted images of identified primary motor neurons and their innervation of epaxial and hypaxial musculature. MiP, middle primary; dRoP, rostral primary, dorsal projection; vRoP, rostral primary, ventral projection; CaP, caudal primary; VaP, variable primary. The distinct types delineate four epaxial and hypaxial muscle groups (color coded and numbered I-IV on right). Dashed line indicating horizontal myoseptum (hm) is provided for reference.

To reveal contribution of individual motor neurons to these nerves and their morphologies, we used a transient, mosaic labeling approach by injecting either mnx:GFP or a combination of mnx:Gal4 or VAChT:Gal4 with UAS:mCD8GFP or UAS:pTagRFP into 1-2 cell stage embryos and performed *in vivo* imaging in 5-7 day old larvae. Traditionally, individual primary motor neurons are identified based on the rostro-caudal soma positions (Myers et al., 1986). Here, we used their peripheral innervation territories in the epaxial and hypaxial muscle to define four distinct quadrants, I-IV (Figure 1e). The ‘middle’ primary (MiP) contributed to the dorsal branch of the ventral root and innervated the entire depth of epaxial muscle in quadrant I via medial nerves. The ‘rostral’ primaries, distinguished by dorsal epaxial or ventral hypaxial innervation, contributed to the ventral branch of the ventral root and innervate the entire depth of quadrants II (dRoP) or III (vRoP) via medial nerves. The ‘caudal’ primary (CaP) also contributed to the ventral branch and innervated the entire depth of quadrant IV via medial nerves. There is also a ‘variable’ primary (VaP) distinguished by its proximity to the CaP (Eisen et al., 1990), that innervated quadrant III.

Next, we used the distinct medial and septal sources of innervation and the delineation of quadrants provided by the primary motor neurons to define different types of secondary motor neurons. One type of secondary could be distinguished based on innervation of axial muscle exclusively via medial nerves (Figure 2a). We named these secondaries ‘m-type’ and identified them according to the quadrants innervated. A second class we named ‘ms-type’, because their axons contribute to both medial and septal nerves (Figure 2b). A third class we named ‘s-type’, because their axons contribute exclusively to the septal nerves (Figure 2c). Among the s-types, we also observed neurons with large, plate-like nerve endings that terminated superficially at the horizontal myoseptum (n = 5) or the dorsal (n = 2) or ventral (n = 2) extremes of vertical myosepta (s’ in Figure 2d,e). Since these neurons appear to be in the process of innervating their respective quadrants, we could not delineate them as we did other s-types, beyond their targeting of the different extremities.

**Figure 2.**
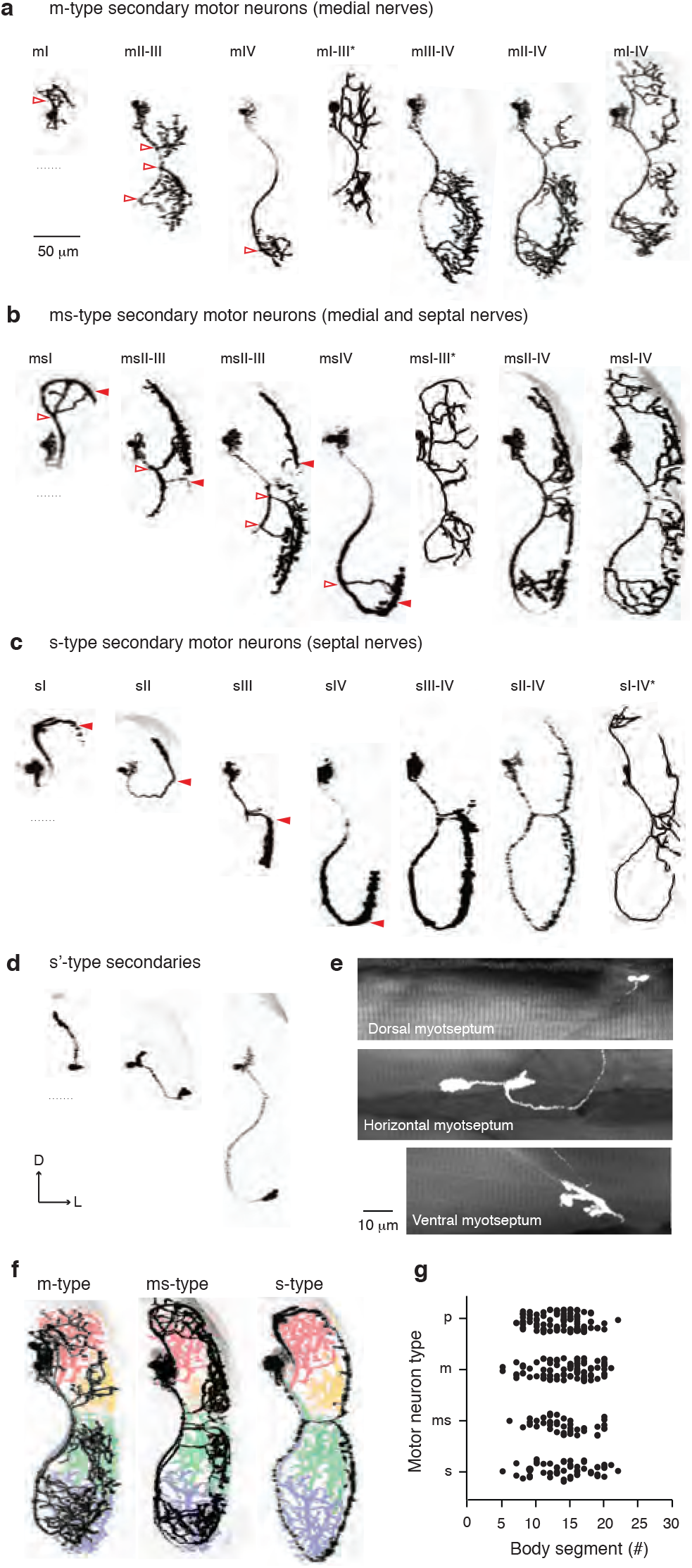
Classifying secondary axial motor neurons based on medial and septal innervation patterns. **a**, Contrast-inverted images of secondary motor neurons in transverse view named based on their contribution to medial peripheral nerves and the epaxial/hypaxial muscle groups they innervate. Asterisks denote reconstructions instead of raw data in cases where one or more motor neurons were also in the field of view. Open red arrowheads illustrate medial nerve projections. **b**, As in *b*, but for secondary motor neurons that contribute to both medial (open red arrowheads) and septal (filled red arrowheads) peripheral nerves. **c**, As in *b*, but for secondary motor neurons that contribute to only septal (red arrowheads) nerves. **d**, As in *b*, but for secondary motor neurons that terminate at the dorsal or ventral extremes of the axial musculature or near the horizontal myoseptum. D, dorsal; L, lateral. **e**, Side on views of the nerve endings of motor neurons shown in d. Collapsed fluorescent images are superimposed on single optical sections of muscle visualized using differential interference contrast. **f**, Examples of secondaries from *b-d* superimposed on color coded primary motor neurons (from Figure 1e) illustrate differences in the medio-lateral distribution of peripheral motor nerve endings between the different types. **g**, Quantification of the rostro-caudal distribution along the body of the different types of motor neuron.

Among the different types a variety of innervation patterns were observed (Table 1), with all m-, ms- and s-type motor neurons innervating a minimum of one quadrant and a maximum of four (Figure 2a-c). Unlike the primaries, there did not appear to be any clear relationship between rostro-caudal soma position and muscle innervation territory among the secondaries. To confirm this, we analyzed soma positions and innervation territories for primaries, which unsurprisingly revealed a significant difference (one-way ANOVA, F(4,90) = 47.8, p < 0.0001). Following post-hoc analysis, of the four primaries, only the dRoP versus vRoP and the CaP versus VaP did not differ significantly in their positions (p > 0.05), as you would expect based on previous work (Eisen et al., 1990; Menelaou and McLean, 2012). Also as expected from previous work (Menelaou and McLean, 2012), there was no significant relationship between rostro-caudal soma position and innervation territory for secondary motor neurons (one-way ANOVA, F(2,182) = 1.3, p = 0.278). This was also true when considering secondaries that exclusively innervated either quadrant I or quadrant IV among the three types (one-way ANOVA, F(5,52) = 0.9, p = 0.459).

**TABLE 1.** Number of observed quadrant innervation patterns

|  | I | II | III | IV | I-II | II-III | III-IV | I-III | II-IV | I-IV | Total (n) |
| --- | --- | --- | --- | --- | --- | --- | --- | --- | --- | --- | --- |
| p-type | 32 | 29 | 27 | 16 |  |  |  |  |  |  | 104 |
| m-type | 6 | - | - | 10 | - | 4 | 2 | 4 | 18 | 53 | 97 |
| ms-type | 8 | - | - | 4 | - | 7 | 1 | 1 | 18 | 3 | 42 |
| s-type | 11 | 1 | 5 | 21 | 1 | 2 | 2 | - | 7 | 1 | 51 |

Compared to the peripheral innervation patterns of primaries, which we will now refer to as ‘p-type’, the termination fields of m-type motor neurons were concentrated in the deeper fast-twitch muscle fibers, s-types were concentrated in the superficial slow muscle fibers, and ms-types innervated intermediate regions containing both fast-twitch and slow fibers (Figure 2f). There also appeared to be very medial fast muscle fibers that were exclusively innervated by p-types, based on gaps in m-type innervation (Figure 2f). Despite differences in the height and width of the axial musculature along the body (Figure 1d), there was no obvious difference in the segmental distribution of each type (Figure 2g).

When viewed from the side, the medial axon of p-type and m-type motor neurons contributed to the dorsal and/or ventral branch of the ventral root near the middle of the muscle segment, from which projections fanned out rostrally and caudally (Figure 3a,b). Multiple, punctate nerve endings were observed terminating diffusely along the full length of the muscle fibers, consistent with patterns of polyneuronal innervation of fast-twitch fibers in teleosts (Johnston, 1981). For ms-type motor neurons, the medial axon followed a similar trajectory and also provided multiterminal innervation of fast-twitch muscle fibers, however the septal branch (or branches) primarily projected rostrally and superficially to innervate the leading edge of the muscle segment (*red arrowheads*, Figure 3c). A subset of ms-types (13 of 42) also innervated the trailing edge of the muscle segment (see also Figure a). The same pattern was true for s-type motor neurons (Figure 3d). Out of 51 s-types, all 51 contributed to septal nerves and innervated the leading edge, while 3 also innervated the trailing edge. Septal branches from ms and s-type neurons ran along the vertical myosepta with relatively little branching. These patterns are consistent with polyneuronal innervation of slow fibers at the myoseptal ends, which represents a more primitive pattern in fishes (Johnston, 1981). Unfortunately, due to the transient nature of labeling, we could not follow these neurons any further in development to see if their axons subsequently extended medially into the muscle segment.

**Figure 3.**
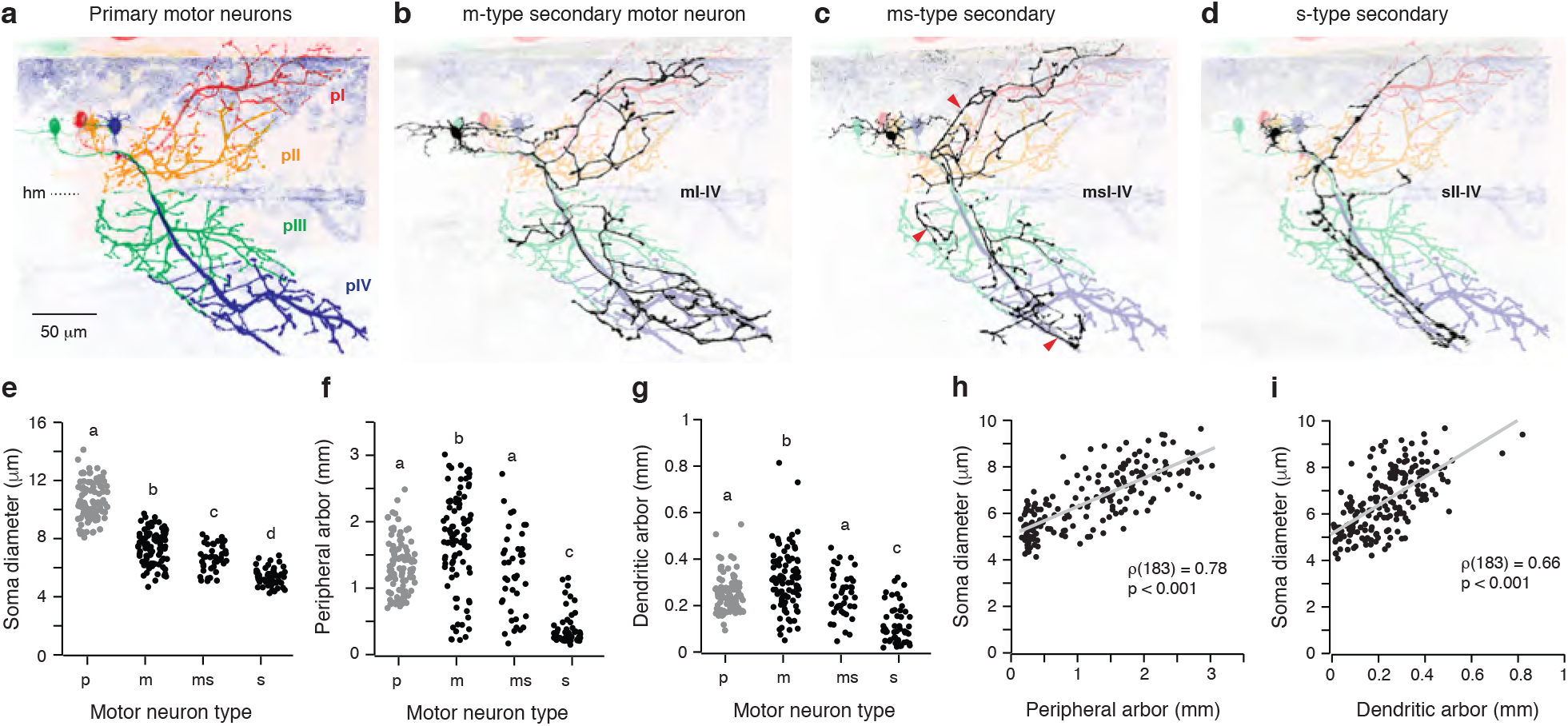
Comparing the morphological features of the different types of axial motor neurons. **a**, p-type motor neurons (pI, pII, pIII, pIV) have been color coded to illustrate their tiling of the dorsal epaxial and ventral hypaxial musculature. Dashed line indicates the horizonal myoseptum (hm). **b**, Example of an m-type motor neuron that innervates all four axial muscle groups (mI-IV) superimposed on p-type motor neurons for reference. **c**, As in b, but an ms-type motor neuron innervating all four muscle groups. Red arrowheads mark septal branches. This is the same neuron illustrated at the far right in Figure 2b. **d**, As in b, but an s-type motor neuron innervating three muscle groups. Note, the intermyotomal cleft runs caudally as you move to the surface of the muscle, so the collapsed image of the s-type peripheral axon is right-shifted relative to the primary reference image. **e**, Quantification of the soma diameter of the different types of motor neuron. Means sharing letters are not significantly different following one-way ANOVA and post-hoc Bonferroni tests. **f**, Quantification of length of the peripheral arbor of different types of motor neuron. Medians sharing letters are not significantly different following Kruskal-Wallis ANOVA and post-hoc Mann-Whitney U tests. **g,** As in *f*, but quantification of the dendritic arbor of different types of motor neuron. **h**, Quantification of soma diameter versus peripheral arbor length of secondary motor neurons. Correlation is reported for Spearman Rank test. **i,** Quantification of the soma diameter versus dendritic arbor length of secondary motor neurons. Correlation is reported for Spearman Rank test.

To further characterize the different types of motor neurons, in a proportion of our dataset (*n* = 280) we analyzed features that would be relevant to their recruitment, specifically soma size, peripheral axon arbor length and dendritic arbor length. As expected, the p-types had the largest somata, with a systematic decrease in soma size from m-type, to ms-type, to s-type motor neurons (Figure 3e). Although there was a clear overlap in values, differences between types were significant (one-way ANOVA, F(3,276) = 314.4, p < 0.001). The m-type and ms-type neurons had a much larger range of peripheral arbor lengths that could exceed that of p-types, consistent with their multi-quadrant innervation patterns (Figure 3f). On the other hand, s-types tended to have smaller peripheral arbors, consistent with their relatively unbranched septal nerve endings (Figure 3f). Statistical analysis revealed a significant difference in peripheral axon length related to type (Kruskal-Wallis ANOVA, H(3) = 120.5, p < 0.001), with m-types representing the largest fields and s-types the smallest following post-hoc analysis (Figure 3f). A similar pattern was observed for dendritic arbor length (Figure 3g). There were significant differences between types (Kruskal-Wallis ANOVA, H(3) = 90.1, p < 0.001), with m-types exhibiting the largest dendritic arborizations and s-types the smallest (Figure 3g).

Among secondaries, a comparison of soma size and either peripheral or dendritic arbor length, revealed a significant positive relationship (Figure 3h,i), where larger neurons tended to have longer axons and dendrites. This observation is also supported by morphological data tabulated according to secondary motor neuron type and number of quadrants innervated (Table 2). A similar, albeit weaker positive correlation between soma size and axon length was found among primaries (Spearman rank correlation, ρ(93) = 0.37, p < 0.001), however there was no significant relationship between soma size and dendrite length (Spearman rank correlation, ρ(93) = 0.20, p = 0.196).

**TABLE 2.** Quantification of morphological features by type and number of quadrants innervated

| | Quadrants innervated | Soma diameter ($\mu\text{m}$) | Dendrite length ($\mu\text{m}$) | Axon length (mm) | Collateral length ($\mu\text{m}$) | Soma DV (normalized) |
| --- | --- | --- | --- | --- | --- | --- |
| m-type | 4 | $7.8 \pm 0.9$ (51) | $354.5 \pm 124.4$ (51) | $2.1 \pm 0.4$ (51) | $131.5 \pm 59.9$ (51) | $0.52 \pm 0.07$ (51) |
| | 3 | $7.5 \pm 1.0$ (19) | $317.7 \pm 102.7$ (19) | $1.8 \pm 0.5$ (19) | $128.6 \pm 52.7$ (18) | $0.54 \pm 0.07$ (19) |
| | 2 | $6.9 \pm 1.3$ (5) | $270.6 \pm 108.8$ (5) | $1.4 \pm 0.3$ (5) | $144.6 \pm 53.1$ (5) | $0.54 \pm 0.11$ (5) |
| | 1 | $5.9 \pm 0.7$ (16) | $163.9 \pm 76.9$ (16) | $0.5 \pm 0.3$ (16) | - | $0.40 \pm 0.07$ (16) |
| ms-type | 4 | $7.3 \pm 0.6$ (3) | $325.3 \pm 74.3$ (3) | $2.1 \pm 0.1$ (3) | $73.7 \pm 30.6$ (3) | $0.46 \pm 0.04$ (3) |
| | 3 | $6.9 \pm 0.7$ (19) | $290.4 \pm 83.1$ (19) | $1.6 \pm 0.4$ (19) | $99.0 \pm 78.3$ (13) | $0.48 \pm 0.07$ (19) |
| | 2 | $6.6 \pm 1.2$ (8) | $238.7 \pm 67.7$ (8) | $1.1 \pm 0.3$ (8) | $93.8 \pm 40.6$ (6) | $0.43 \pm 0.06$ (8) |
| | 1 | $5.8 \pm 0.6$ (11) | $144.0 \pm 66.2$ (11) | $0.4 \pm 0.2$ (11) | - | $0.41 \pm 0.07$ (11) |
| s-type | 4 | 5.9 (1) | 243 (1) | 1.1 (1) | 21.6 (1) | 0.51 (1) |
| | 3 | $5.4 \pm 0.8$ (5) | $222.6 \pm 93.6$ (5) | $0.9 \pm 0.2$ (5) | $18.9 \pm 0.2$ (2) | $0.44 \pm 0.11$ (5) |
| | 2 | $5.7 \pm 0.5$ (5) | $122.8 \pm 67.4$ (5) | $0.6 \pm 0.2$ (5) | - | $0.46 \pm 0.07$ (5) |
| | 1 | $5.1 \pm 0.6$ (33) | $96.1 \pm 76.1$ (33) | $0.3 \pm 0.1$ (33) | - | $0.29 \pm 0.10$ (33) |
| s'-type | - | $5.4 \pm 0.4$ (9) | $70.1 \pm 34.7$ (9) | $0.2 \pm 0.1$ (9) | | $0.23 \pm 0.09$ (9) |

Data reported as mean ± standard deviation. n's are reported parenthetically. Note, for some neurons we did not collect higher resolution images (9 of 104 p-types, 6 of 97 m-types, 1 of 42 ms-types, and 7 of 51 s-types; total 23 of 303). DV, dorso-ventral position.

These data suggest that axial motor units in larval zebrafish divide up fast and/or slow muscle into four distinct quadrants, with units as small as one quadrant and as large as four. Our data also confirm the existence of three distinct anatomical types of secondary motor neuron, defined by their size and target musculature (Menelaou and McLean, 2012; Asakawa et al., 2013); large m-units that innervate fast-twitch muscle fibers, small s-units that innervate slow fibers, and intermediate ms-units that innervate fast-twitch and slow fibers. The overlapping patterns of innervation observed within and between types is consistent with polyneuronal innervation of muscle fibers. Thus, distinct motor units in larval zebrafish are comprised of different motor neurons that target common fast-twitch and/or slow muscle fibers.

### Central axon collaterals are observed primarily in larger fast-twitch units

Next, we asked how central axon projections relate to the different types of axial motor unit. In zebrafish, axial motor pools are strictly segmental with axons exiting via the ventral root situated at the caudal edge of the pool (Figure 4a). Only rarely (n = 2 of 303), did we did find motor neurons whose axons exited two consecutive roots (data not shown) and these were both ms-types. In every type of motor neuron, we found instances where collaterals from the main axon could be observed prior to exiting the ventral root (Figure 4b,c). Axon collaterals arose from single or multiple processes and varied in their extent of branching among primaries (Figure 4b) and secondaries (Figure 4c). Axon collaterals terminated locally and could project dorsally up to the level of the soma of origin, rostrally beyond the soma of origin and caudally into the next spinal segment (Figure 4b,c).

**Figure 4.**
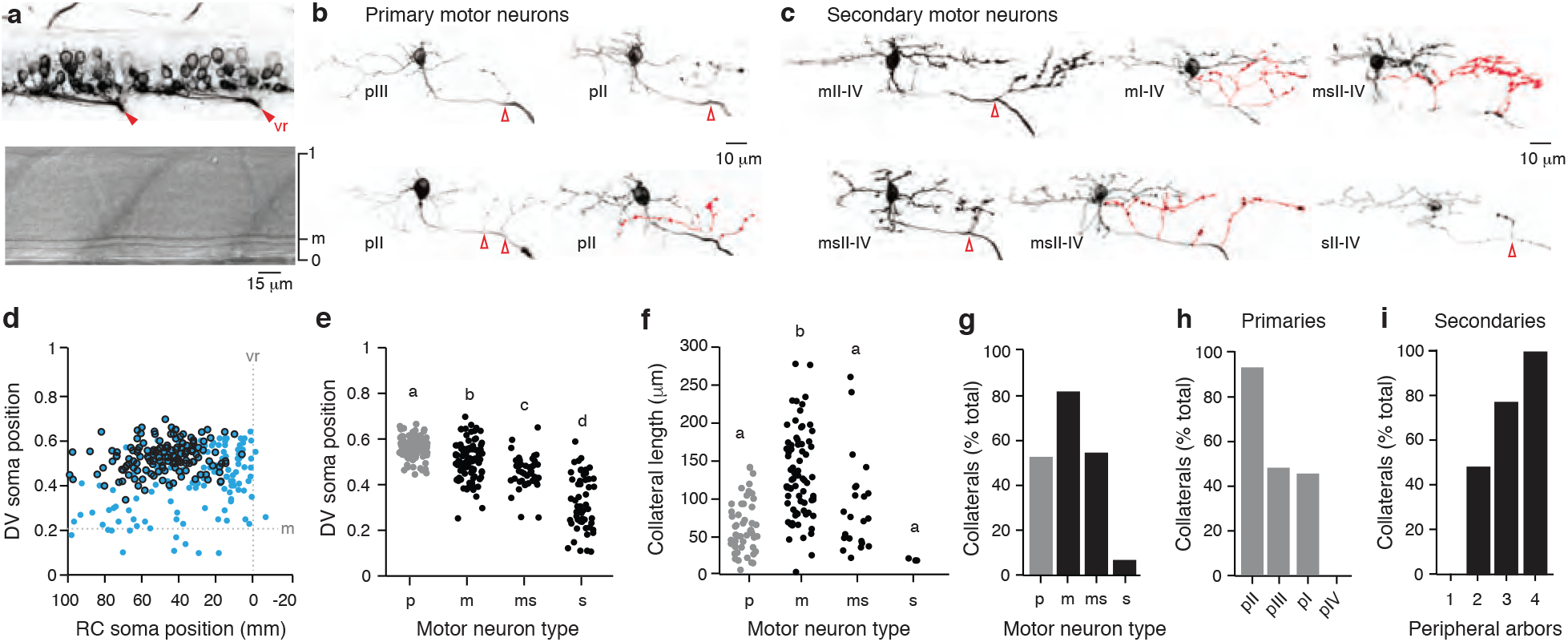
Characterizing motor neurons with central axon collaterals. **a**, Top: contrast-inverted image of spinal motor neurons in Tg[mnx1:GFP] fish viewed from the side. The ventral roots (vr) are indicated by red filled arrowheads. Bottom: differential interference contrast image of the same location in spinal cord shown above illustrates major landmarks for normalization (1, dorsal edge of spinal cord, 0, ventral edge of spinal cord, m, Mauthner axon). **b**, Contrast-inverted images of p-type motor neurons. Central axon collaterals are highlighted in red or with red open arrowheads. **c**, As in *b*, but for m-, ms-, and s-type motor neurons. **d**, Quantification of the dorso-ventral position of motor neuron somata normalized to the height of the spinal cord and their distance from the ventral root exit point. Blue indicates total dataset, while black indicates position of motor neurons with central axon collaterals. **e**, Quantification of normalized dorso-ventral soma positions of the different types of motor neuron. Means sharing letters are not significantly different following one-way ANOVA and post-hoc Bonferroni tests. **f,** Quantification of the central collateral length of different types of motor neuron. Means sharing letters are not significantly different following one-way ANOVA and post-hoc Bonferroni tests. **g**, Quantification of likelihood of central collateral among the different types of motor neuron. **h,** Quantification of the likelihood of central collaterals among p-type motor neurons, ordered from left to right by mean rostro-caudal soma position, pII = 42 ± 12 μm, pIII = 41 ± 21 μm, pI = 31 ± 10 μm, pIV = 8 ± 4 μm. Note that pIIIs contain VaPs = 13.03 ± 8 μm and vRoPs = 51.39 ± 12 μm. Since VaPs never exhibit central collaterals, this brings down the average. **i,** Quantification of central collateral presence in secondary motor neurons based on the number of axial muscle quadrants innervated.

Central axon collaterals were found in only half of the dataset (n = 49 of 95 primaries, n = 99 of 185 secondaries), which tended to be located more rostrally and more dorsally in the spinal segment (Figure 4d), consistent with our previous study (Menelaou and McLean, 2012). Since motor neurons located more dorsally are active at faster speeds (McLean et al., 2007; Knafo et al., 2017), we compared the relative dorsoventral location of the different types of motor neurons. Consistent with the size, muscle targets, and recruitment order, p-types were the most dorsal, followed by m-types, then ms-types and finally s-types which were distributed most ventrally (Figure 4e). Notably, there was some intermingling between types, meaning that the discrete types are not clearly segregated into pools as motor neurons appear to be in adult zebrafish (Ampatzis et al., 2013). Nevertheless, following statistical analysis, these differences proved significant (one-way ANOVA, F(3,276) = 106.1, p < 0.001).

To further characterize axon collaterals, we analyzed their length and likelihood of occurrence among the four types of motor neurons. The m-type and ms-type neurons have a larger range of collateral lengths that can exceed that of p-types, while s-type neurons have the smallest collateral lengths (Figure 4f; Table 2). Statistical analysis demonstrated a significant difference (one-way ANOVA, F(3,144) = 20.2), which post-hoc analysis revealed was due to m-types having the longest collaterals compared to the other types (Figure 4f). When we considered the overall likelihood of collaterals occurring, they were most likely in m-type neurons, with p-type and ms-type neurons having about the same likelihood (Figure 4g). On the other hand, s-type neurons rarely exhibited collaterals (Figure 4g). Among primaries, cells located more rostrally have a higher likelihood of collaterals than more caudal neurons (Figure 4h). This meant that RoPs and MiPs were more likely to exhibit collaterals, while CaPs and VaPs never did. Among secondaries, those that exhibit a greater number of innervated quadrants had a higher likelihood of exhibiting collaterals (Figure 4i). For the rare cases of collaterals in s-types, these were either sI-IV (n = 1) or sII-IV neurons (n = 2). Thus, among secondaries, the likelihood of having collaterals is not only related to being sufficiently rostral enough in the segment to have the opportunity to extend axon collaterals before exiting the ventral root, but also how much muscle they innervate after the exit.

Next, to confirm axon collaterals were output and not input structures, we injected a combination of mnx:Gal4 or VAChT:Gal4 with UAS:GFP or UAS:pTagRFP to visualize motor neuron morphology and either UAS:syp-pTagRFP or UAS:syp-GFP to label collaterals with the presynaptic marker synaptophysin or UAS:PSDGFP to label collaterals with the postsynaptic marker PSD95 (Figure 5a). For this analysis, we pooled the primary and secondary populations. Punctate synaptophysin labeling was observed out in the periphery and confirmed different medio-lateral distributions of muscles targeted by the different types (Figure 5b). Furthermore, there was a good correspondence between the number of peripheral synaptophysin puncta and peripheral axon length (Figure 5d), confirming the utility of arbor length as a measure of functional output.

**Figure 5.**
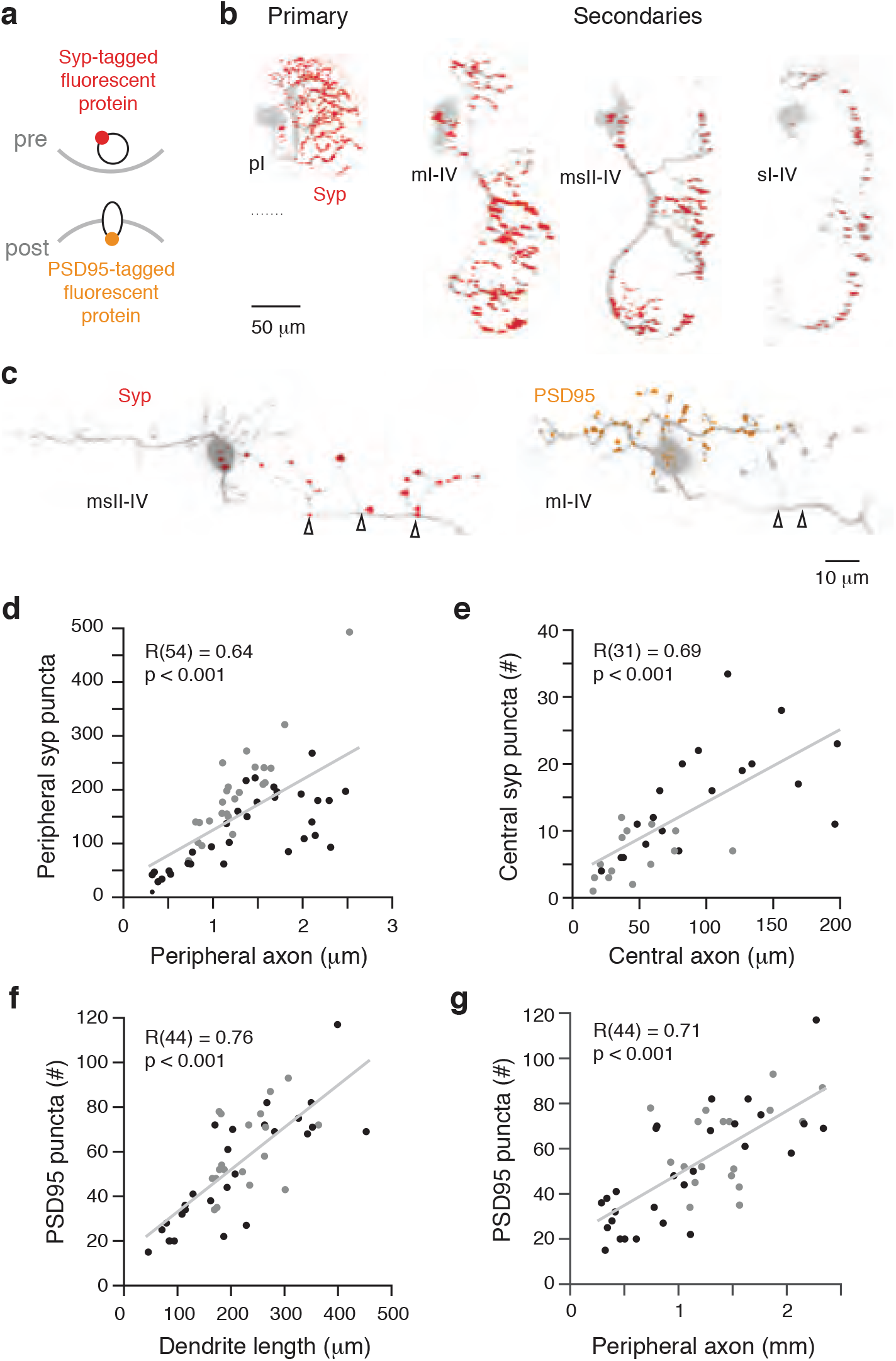
Characterizing the input/output features of the different types of motor neurons. **a**, Schematic illustrating the strategy for identifying pre- and post-synaptic structures in motor neurons using fluorescently tagged synaptophysin (red, pre-synaptic), and PSD95 (orange, post-synaptic). **b**, Contrast-inverted images of motor neurons (grey) in transverse view from each type illustrating synaptophysin (Syp) expression (red). **c**, Contrast-inverted images of motor neurons from the side view illustrating synaptophysin labeling on axon collaterals (left) and PSD95 labeling on dendrites and soma (right). Open black arrowheads demarcate central axon collaterals. **d**, Quantification of motor neuron peripheral synaptophysin puncta number versus peripheral axon length. Secondary motor neurons labeled in black and primary motor neurons in grey. Correlation is reported for Pearson’s test. **e**, As in *d*, but quantification of motor neuron central synaptophysin puncta number versus central collateral length. **f**, As in *d*, but quantification of PSD95 puncta number versus dendrite length. **g,** As in *d*, but quantification of PSD95 puncta number versus peripheral axon length. Correlation is reported for Pearson’s test.

Imaging within spinal cord revealed punctate synaptophysin expression distributed throughout the axon collaterals that was missing in the dendrites (Figure 5c, *left*). This contrasts the expression pattern of PSD95, which was distributed throughout the dendrites, but missing in the collaterals (Figure 5c, *right*). Unfortunately, due to the aggregation of PSD95 fluorescence in the soma, we could not unambiguously identify somatic contacts, however they did not appear very abundant. As with the peripheral innervation patterns, there was a good correspondence between synaptophysin puncta number and central axon length (Figure 5e). Similarly, PSD95 puncta number and dendrite length were positively correlated (Figure 5f). A comparison of PSD95 puncta number and peripheral axon length also revealed a clear input-output relationship, where motor neurons with more output receive more input (Figure 5g).

These data demonstrate that axon collaterals are more likely observed in fast motor units, which is consistent with previous work in cats and mice (Cullheim and Kellerth, 1978; Friedman et al., 1981; Bhumbra and Beato, 2018). What we also reveal is that collaterals are also more frequently observed in neurons with larger innervation territories, regardless of unit identity. In addition, our findings also demonstrate a clear input-output function, where the levels of excitatory afferent drive to motor neurons are scaled to the levels of their peripheral and central innervation. This is consistent with voltage-clamp recordings that demonstrate larger, lower Rin motor neurons receive greater amounts of excitatory drive (Kishore et al., 2014) and also with electron microscopy data demonstrating smaller motor neurons receive less synaptic input (Svara et al., 2018).

### Central axon collaterals ramify in the neuropil with potential perisomatic motor neuron connections

Our next goal was to identify the potential targets for axon collaterals within the spinal cord. In the transverse plane, axial motor neurons form a column in the ventral half of spinal cord within a medial soma layer flanked by lateral neuropil layers (Figure 6a). The original description of primary and secondary motor neurons in larval zebrafish distinguished them based on their size and their axon trajectories relative to the giant Mauthner axon (Myers, 1985), which is thought to reflect axo-axonic connections for powerful escape maneuvers (Fetcho and Faber, 1988). Specifically, primary axons passed under the Mauthner cell axon, while secondary axons passed over it. Among our own dataset (Figure 6b), we also found that all p-types passed under the Mauthner axon (n = 95 of 95). However, the picture for secondaries was more nuanced. Among m-types, 88 out of 91 passed under (∼97%), for ms-types it was 34 out of 41 (∼83%), for s-types it was only 24 out of 44 (∼55%), and for s’-types we never observed the axon passing under the Mauthner axon (0%; n = 9). The transverse view also revealed a good correspondence between the dorso-ventral distribution of dendrites and somata (Figure 6b). To see if this pattern was a systematic one, we plotted dorsoventral soma position against the dorsal-most extent of dendrites, which revealed a significant positive correlation (Figure 6c, *left*). A similar relationship was not found when considering the dorsal-most extent of axon collaterals (Figure 6c, *right*), which is to be expected since motor neurons with collaterals are concentrated in more dorsal regions of cord (see Figure 4d).

**Figure 6.**
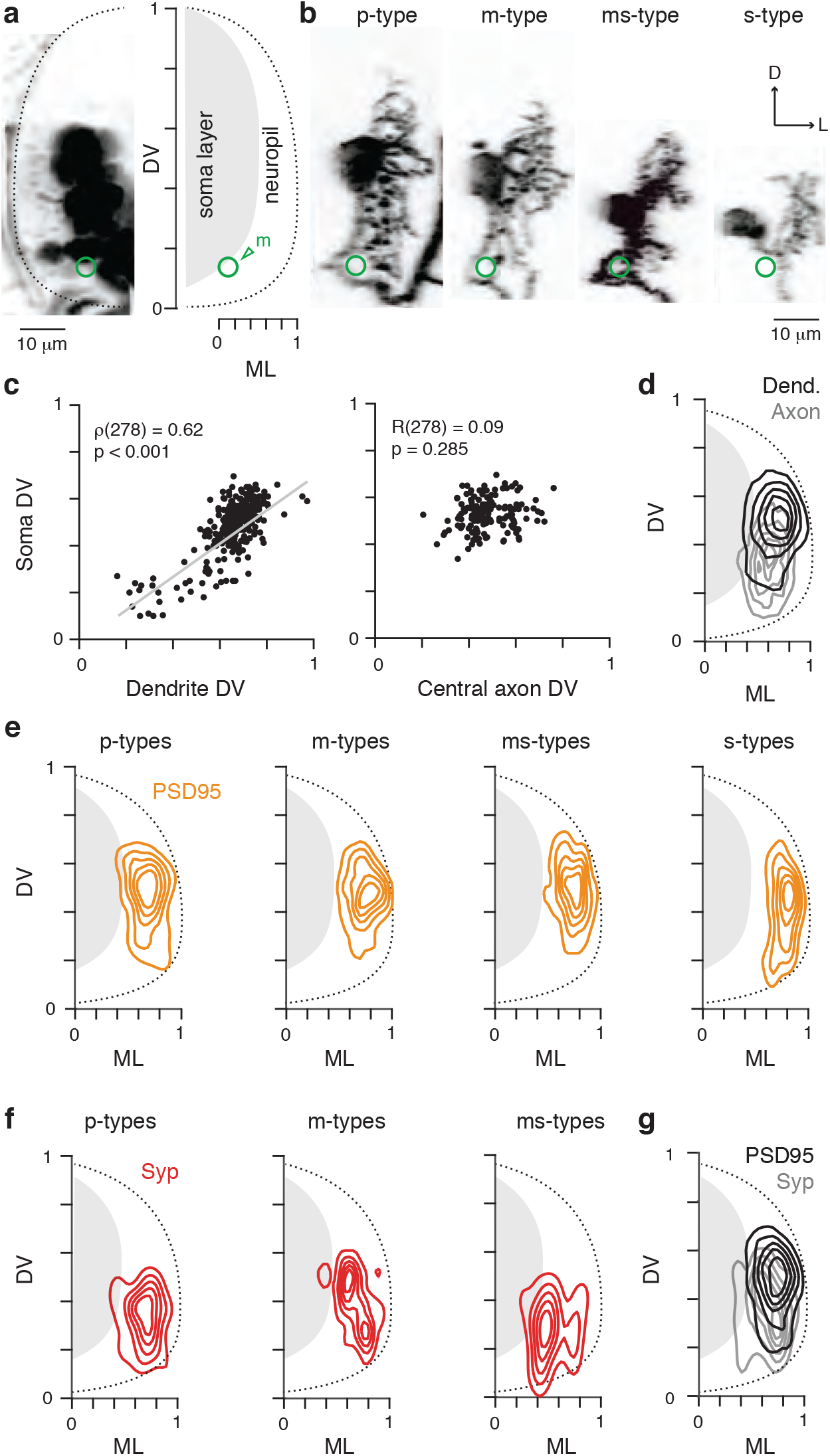
Assessing the topographic organization of the input/output of the different types of motor neurons. **a**, *Left*: A confocal z-stack in transverse view illustrating the medio-lateral (ML) and dorso-ventral (DV) distribution of motor neuron somata and neuropil within the spinal cord in a Tg[mnx1-3×125bp:Gal4-VP16;UAS:pTagRFP] fish. *Right*: Schematic demonstrating the normalization according to anatomical landmarks, which include the boundary of spinal cord (dashed lines) and the Mauthner axon (green, m). **b,** Depiction of axon trajectory and proximity to the Mauthner axon (green, m) in transverse view of the different types of motor neuron prior to exiting the spinal cord. D, dorsal; L, lateral. **c**, *Left*: Quantification of the maximum normalized dendrite dorso-ventral (DV) height versus the normalized soma DV position. Correlation is reported for Spearman Rank test. *Right*: Quantification of the maximum normalized central collateral DV height versus the normalized soma DV position. Correlation is reported for Pearson’s test. **d**, Contour density plot of the normalized medio-lateral and dorso-ventral distribution of the dendrites (black) and central collaterals (grey) of motor neurons with central collaterals (n = 148). The innermost contour represents the highest filament density. **e,** Contour density plot of the normalized medio-lateral and dorso-ventral distribution of PSD95 puncta of the different types of motor neuron (p-type, n = 19; m-type, n = 10; ms-type, n = 11; s-type, n = 6). The innermost contour represents the highest puncta density. **f**, As in *d*, but contour density plot of the normalized medio-lateral and dorso-ventral distribution of synaptophysin puncta of the different types of motor neuron (p-type, n =15; m-type, n = 12; ms-type, n = 5). **g**, As in *d*, but contour density plot of the normalized medio-lateral and dorso-ventral distribution of synaptophysin and PSD95 puncta from all the types of motor neuron (n = 46 motor neurons with PSD95 labeling and n = 32 motor neurons with synaptophysin labeling).

To get a better sense of the dorso-ventral and medio-lateral distribution of dendrites and collaterals, we used contour density plots of filament lengths normalized to anatomical landmarks (Figure 6a). Axon collaterals tended to be more ventrally distributed compared to dendrites, but both were concentrated laterally in the neuropil (Figure 6d). We then linked these input-output patterns to more concrete representations by plotting the contour densities of PSD95 and synaptophysin puncta according to motor unit type. PSD95 distribution was similar for p-types, m-types and ms-types, but tended to be distributed more laterally and ventrally for s-types (Figure 6e). Synaptophysin distribution was more heterogeneous (Figure 6f). In p-types putative outputs were concentrated ventrally. For m-types, outputs were concentrated dorsally and ventrally in the neuropil, while for ms-types they were concentrated more ventrally and medially in the neuropil. Unfortunately, we did not have sufficient numbers of s-types to perform a similar output analysis, since they rarely exhibited collaterals. Regardless, by pooling all the PSD95 and synaptophysin data across types, a similar pattern to the dendrite and collateral filament analysis was observed (Figure 6g).

Because the majority of axon collaterals were concentrated in the neuropil, it was difficult to identify putative synaptic contacts based on proximity. Also, the intermingling of dendritic territories among types prevented any potential link to unit identity based on neuropil distribution. However, since many of our injections to label individual motor neurons were performed in Tg[mnx1:GFP] or Tg[mnx1:Gal4;UAS-pTagRFP] lines (n = 75 of 303), in a proportion of these experiments (n = 17) we observed central nerve terminals in close proximity to other motor neuron somata (Figure a). Since the filament and synaptophysin analysis were in good correspondence (cf., Figure 6d,g), these likely reflect synaptic connections. An analysis of the dorso-ventral soma locations of putative target neurons pattern was also consistent with synaptophysin contour density analysis, where p-types targeted motor neurons below their own positions, while m-types and ms-types could target motor neurons located above and below them (Figure 7b). In addition, on one occasion we labeled two motor neurons with a single injection (an m-type neuron and an ms-type neuron innervating the same quadrants but in neighboring segments), which allowed us to visualize their morphologies and synaptic output (Figure 7b). The axon collateral and synaptophysin puncta from the m-type in the same optical section (0.7 micrometer) as the proximal dendrite and soma of the ms-type (Figure 7c).

**Figure 7.**
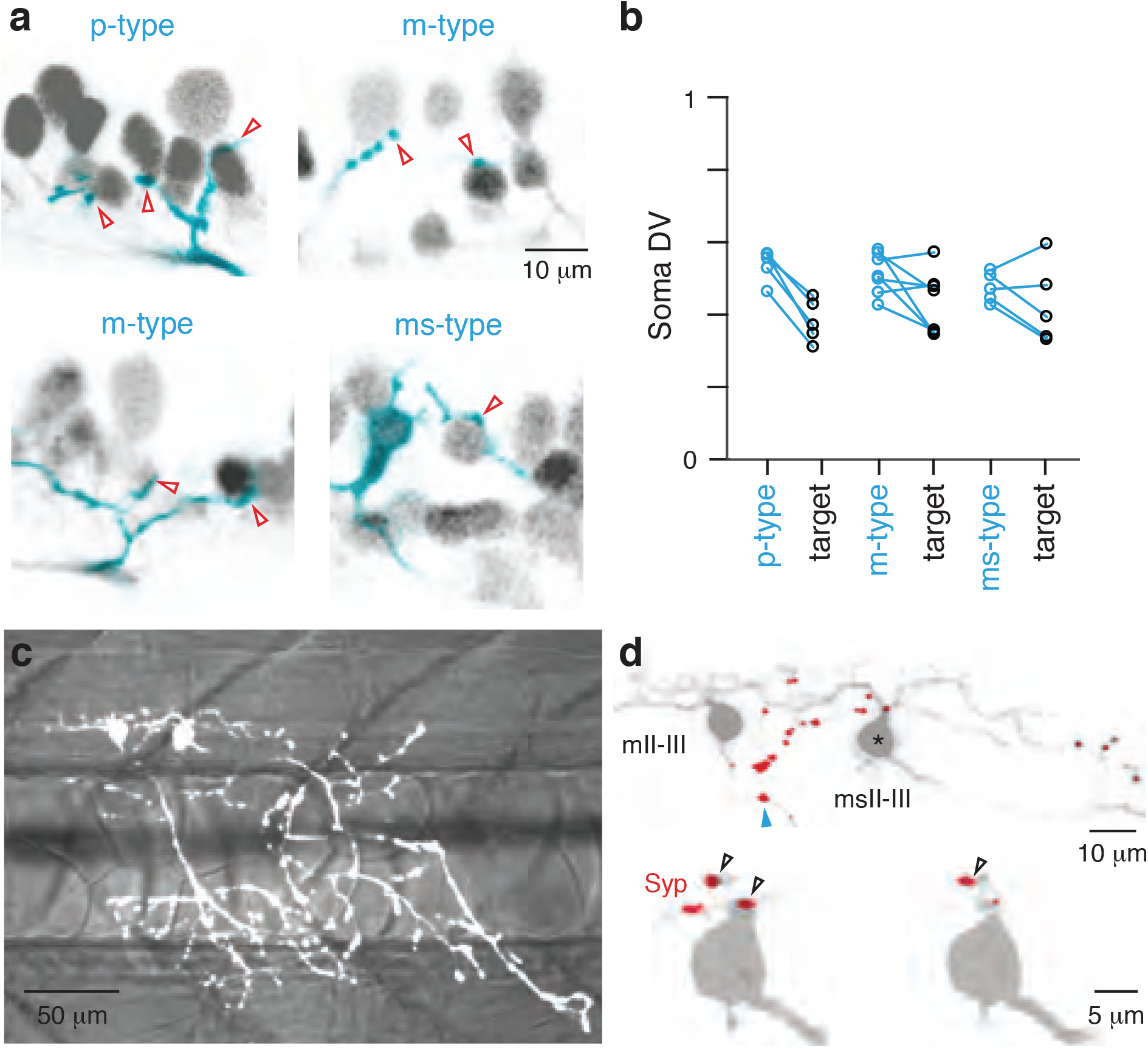
Identifying putative motor neuron-motor neuron contacts. **a**, Single optical sections viewed from the side of putative contacts (open red arrowheads) between individual motor neurons of different types labeled by injections and all axial motor neurons in the Tg[mnx1-3×125bp:Gal4-VP16;UAS:pTagRFP] line. Dorsal is up and rostral is left. **b**, Quantification of dorso-ventral location of p-(n = 5), m-(n = 7), and ms-type (n = 5) motor neurons and their putative motor neuron targets. **c**, Collapsed fluorescent image of an mII-III and msII-III secondary motor neuron in muscle segments 16 and 17, respectively, superimposed on a differential interference contrast (DIC) image to illustrate segment boundaries. **d**, *Top*: Contrasted-inverted image of motor neurons from *a* in grey and synaptophysin puncta labeling in red. Blue arrowhead marks central collateral from rostral m-type neuron. *Bottom*: Single optical sections of neuron labeled with asterisk above, depicting putative perisomatic connectivity from the mII-III to the msII-III neuron (open black arrowheads).

Collectively, the distribution of synaptophysin in axon collaterals and their proximity to other motor neurons is consistent with chemical synaptic transmission between motor neuron types, which could be cholinergic and/or glutamatergic (Perrins and Roberts, 1995; Bhumbra and Beato, 2018). The p-types almost certainly contact motor neurons outside their type, given the ventral distribution of their outputs. A similar hierarchical arrangement is observed for m-types, which can innervate ms-types even in neighboring segments where they presumably do not innervate the same muscle fibers. For m-types, there is evidence for central nerve terminals near the somata of p-types, which would suggest that secondaries can also excite primaries. While the dorso-ventral distribution of dendrites matches dorso-ventral soma distributions, there is sufficient variability in positioning related to type that no systematic relationship between dendritic arborization fields and type was observed. Since axon collaterals are concentrated in the neuropil, interactions within and between motor neuron types and other interneurons are possible but will need to be confirmed.

## 4. Discussion

Our goal was to define types of axial motor units in larval zebrafish and to assess their central axon collaterals and putative targets in the spinal cord to provide a foundation for future functional studies leveraging the experimental advantages of this model system. By using the tiled innervation patterns of primary motor neurons to define four epaxial and hypaxial muscle groups (Figure 8a) and by comparing the relative medio-lateral distribution of axonal arbors with confocal microscopy, we now have a clearer picture of the unit organization of secondary motor neurons and their central and peripheral innervation patterns.

**Figure 8.**
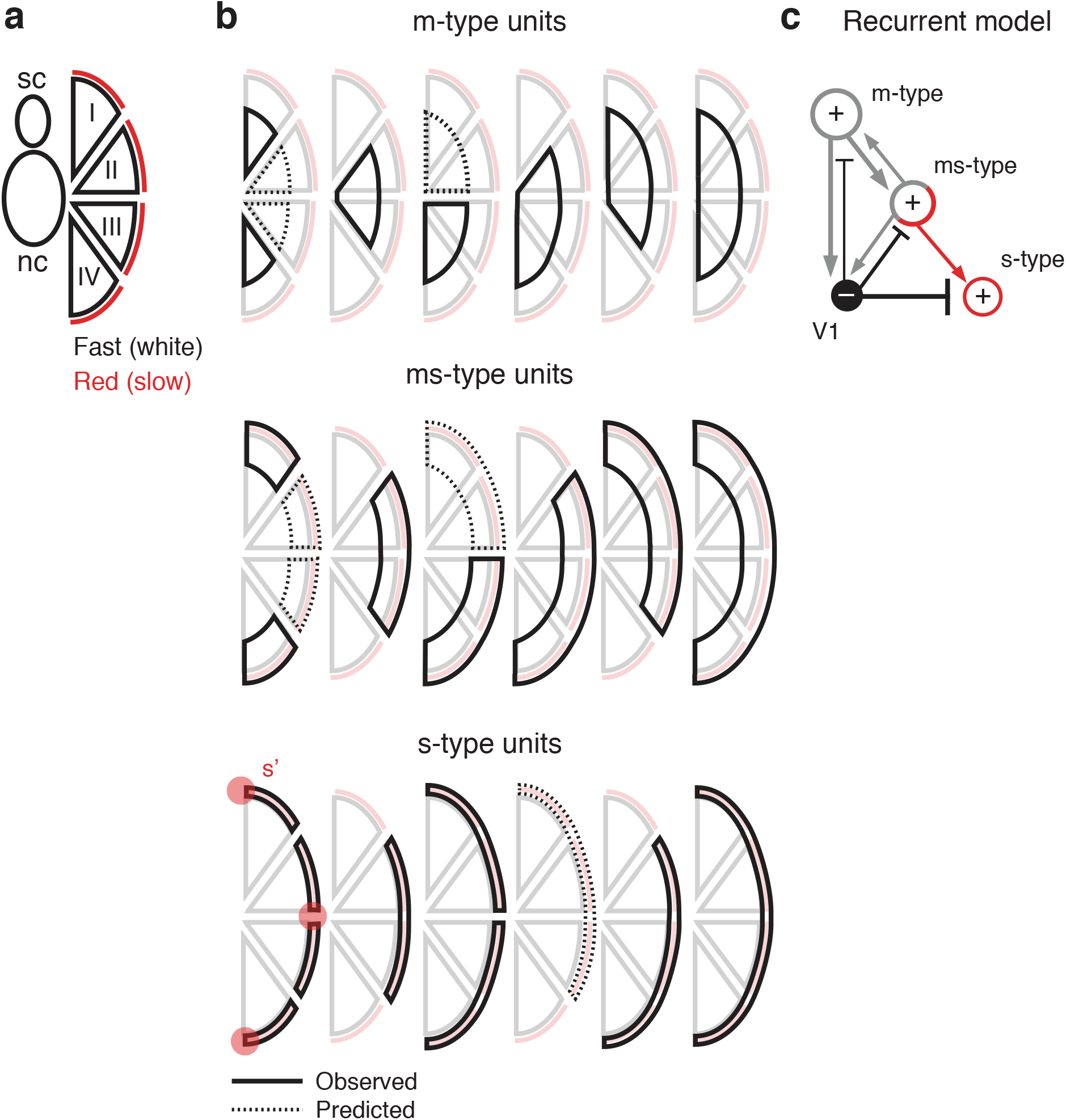
Summary of the main observations. **a**, Schematic depicting the major divisions of the axial muscle based on primary motor neuron innervation. sc, spinal cord; nc, notochord. **b**, Schematics depicting the proposed muscle innervation territories of the newly characterized motor neuron types as they transition from deep-fast innervating muscle to superficial-slow innervating muscle (top to bottom). Immature s-types (s’) terminate in the extremities noted by red circles. Territories are superimposed on primary quadrants. **c,** Schematic depicting the proposed recurrent circuitry involving motor neurons defined by peripheral innervation patterns. Fast-twitch units (m-type and ms-type) are more likely to have collaterals and be interconnected. We also propose fast units would be more likely to innervate the equivalent of Renshaw cells in fish, namely V1 interneurons (see Discussion for details), which have more potent inhibitory inputs to slow units.

We find that secondary motor neurons can be divided into three distinct units (m-type, ms-type, s-type) based on muscle targeting with unit sizes ranging from one quadrant up to four (Figure 8b). The combinations of quadrants innervated was highly stereotyped from fish to fish and tended to be contiguous and not discrete. For example, out of the 303 motoneurons in our dataset, we never observed motor neurons with a gap between their innervated quadrants (e.g., innervating quadrant I and IV). Indeed, the stereotypy we observe related to innervation patterns allows us predict some motor neurons that may exist but were not labeled here. For example, among m- and ms-types, we never observed motor neurons innervating only quadrant II and quadrant III or neurons innervating both quadrant I and quadrant II (Figure 8b). Given the probabilistic nature of our labeling method and the greater likelihood of labeling earlier born neurons (cf., Table 1), we cannot rule out that these missing types of units exist but are rare or develop later on. Regardless, there are numerous combinations in which units could be deployed to provide graded postural control over a range of forces and speeds (Bagnall and Schoppik, 2018).

Previous work has divided secondary motor neurons into three broad types based on the trajectory of their main axon (Menelaou and McLean, 2012; Asakawa et al., 2013). Under our new classification scheme, bifurcating, ventral and dorsal secondaries would now be called ‘m-type’, intermyotomal secondaries with collaterals would be ‘ms-type’ and intermyotomal secondaries with no collaterals ‘s-type’. Moreover, the distribution of different types along the dorso-ventral axis is consistent the topographic pattern of recruitment at this age (McLean et al., 2007), although there is some intermingling among the different types. Consequently, the links between electrophysiological properties and muscle fiber innervation described using previous anatomical designations would be expected to hold with our new classifications. Thus, more persistent activation of slow muscle fibers is achieved by smaller, lower threshold s-type motor neurons with intrinsic burst firing properties and weaker, but potentiating neuromuscular synapses, while transient activation of fast muscle fibers is achieved by larger, higher threshold m-type motor neurons with chattering firing properties and stronger, but more depressing neuromuscular synapses (Menelaou and McLean, 2012; Wang and Brehm, 2017).

A recent electron microscopy study has also defined three groups of unmyelinated secondary motor neurons based on their size and their initial axon trajectories as they exit the ventral root (Svara et al., 2018). ‘Small’ and ‘medium’ motor neurons were distinguished by contributions to either the dorsal or ventral branch of the ventral root. These likely include the smaller s-type, ms-type and m-type motor neurons with more restricted epaxial or hypaxial innervation territories. ‘Large’ neurons contributed to the both the dorsal and ventral branches and so likely include larger m-type and ms-type motor neurons with multiple epaxial and hypaxial innervation territories. Unfortunately, since full axon reconstructions were not possible given the volume of their dataset, it is difficult to definitively link these classifications to ours, particularly given the partial overlap in soma size between m-, ms- and s-type units depending on the number of quadrants innervated (Table 2). Also, since our assessment was over multiple segments, we cannot say for sure whether all of the types we report here are present in a single hemi-segment. However, given that there are reported to be just over 60 motor neurons per hemi-segment it is certainly possible (Asakawa et al., 2013; Svara et al., 2018).

The same study also revealed small motor neurons that lack dendrites and chemical synaptic inputs, suggesting that they are not yet integrated into the locomotor network (Svara et al., 2018). We did not observe any s-types that lacked dendrites, however, this could be because adendritic neurons are relatively rare (< 9% of the total population; Asakawa et al., 2013; Svara et al., 2018) and immature, which would make them harder to find using our mosaic labeling strategy. However, we did observe s-type neurons whose axons appeared to be arrested at the horizontal myoseptum and dorsal and ventral extremes of the vertical myoseptum (s’-types). Given that s’-types do have dendrites, albeit the shortest ones (Table 2), these neurons could provide extremely fine control of slow muscle at the extremities. Alternatively, it could be that these relatively immature neurons are also waiting for their target muscles to differentiate or mature, since the regions targeted by s’-types are sources of new muscle fibers as larvae grow into adults (Hollway and Currie, 2003; Gurevich et al., 2015). Unfortunately, due to the transient nature of our labeling method, we could not confirm the ultimate target of s’-types, whether new axial or fin musculature (Siomava et al., 2018).

Our work also suggests that at larval stages the solution to generating intermediate forces and speeds is to innervate more lateral fast-twitch and slow muscle fibers via ms-type secondary motor neurons. By adulthood, zebrafish have developed axial muscle fibers with intermediate ‘pink’ contractile properties that can be found in the transition from slow to fast-twitch muscle fibers at the horizontal myoseptum and also interspersed among the fast-twitch epaxial and hypaxial muscle at their dorsal and ventral extremes (van Raamsdonk et al., 1982). These are regions innervated by ms-type neurons in larvae. Also, motor neurons in adult zebrafish targeting superficial slow muscle, intermediate pink muscle and deeper fast-twitch muscle are recruited during fictive swimming in an orderly and topographic fashion from slow-intermediate-fast (Ampatzis et al., 2013). While it remains to be seen if adult fast motor neurons are m-type, intermediate motor neurons are ms-type and slow are s-type, the morphological and electrophysiological properties exhibited by fast, intermediate and slow neurons are at least consistent with our classifications (Ampatzis et al., 2013). Studies of the development of muscle fibers from larvae to adults have revealed evidence for both stratified and mosaic hyperplasia (Barresi et al., 2001; Patterson et al., 2008) and so intermediate fibers could be new fibers with new metabolic properties or existing fibers that acquire new properties after their fusion with new fibers.

Critically, the definition of distinct units based on innervation patterns allowed us to link the presence or absence of central collaterals to functional output. Among primaries, the more rostrally positioned neurons within the segment are more likely to exhibit collaterals, meaning drive to medial and dorsal fast-twitch muscles is being fed back into spinal cord. Among secondaries, motor neurons innervating fast-twitch muscles are more likely to exhibit axon collaterals with increasing likelihood related to number of quadrants innervated. Since we also find that larger units with more central and peripheral output also receive more putative excitatory synaptic contacts, this suggests that during increases in swimming speed, the drive responsible for recruiting larger units is being fed back into spinal cord. Whatever the purpose, it is unlikely to be related to pool-specific commands, since secondaries innervating both epaxial and hypaxial muscle are more likely to exhibit collaterals. Because central collaterals are concentrated in the neuropil we could not easily assess putative targets based on proximity, however we did find evidence for motor neuron-motor neuron connections within and between segments. At least for the one example we provide, motor neurons with intersegmental connections appear to innervate the same quadrants. Consequently, the simplest explanation that can account for our anatomical observations is that motor neurons innervating synergistic groups of muscle are also interconnected via central collaterals. In this scenario, motor neurons innervating four quadrants would exhibit more axon collaterals because there are more motor units that share target musculature, followed by neurons innervating three, then two, and so forth. If this is true, it would suggest that one of the major functions of central collaterals is to reinforce patterns of peripheral activation during movements of increasing force and speed.

Our observation that recurrent central collaterals are more likely to be found in earlier born fast motor units is consistent with physiological studies in frog embryos, neonatal/adult mice and adult cats (Cullheim and Kellerth, 1978; Perrins and Roberts, 1995; Bhumbra and Beato, 2018). Physiological studies have also revealed motor neuron connections to inhibitory Renshaw cells, which provide recurrent inhibitory feedback (Renshaw, 1941; Eccles et al., 1954; Nishimaru et al., 2005; Bhumbra et al., 2014; Moore et al., 2015; d’Incamps et al., 2017). In cats, it has been reported that slower motor units are subject to stronger Renshaw-mediated inhibition, which leads to their de-recruitment during strong movements, presumably to prevent overstimulation of slow muscle fibers (Friedman et al., 1981). It has also been reported in cats that Renshaw cells primarily target motor neuron dendrites (Fyffe, 1991). Since our data reveal that motor neuron central collaterals are also predominantly distributed in the neuropil, these observations collectively suggest that whatever computation is being performed by recurrent circuitry, it is occurring out in the dendrites.

In larval zebrafish, slower motor neurons receive stronger inhibition concurrent with excitation than faster motor neurons during high frequencies of ‘fictive’ swimming (Kishore et al., 2014), which can lead to them being silenced (Menelaou and McLean, 2012). The same is also true in adult zebrafish, where slow motor neurons are inhibited during high frequency ‘fictive’ escape responses (Kyriakatos et al., 2011; Song et al., 2015). A potential source for this inhibition is ipsilaterally-projecting En1-labeled glycinergic interneurons derived from the p1 domain (V1 neurons), from which Renshaw cells are derived (Higashijima et al., 2004; Li et al., 2004; Benito-Gonzalez and Alvarez, 2012; Stam et al., 2012). Consequently, a prediction worth testing in light of our findings is that recurrent collaterals not only support the activation of synergistic fast units during strong movements, but also the concurrent inhibition of slow units via V1 inhibitory interneurons (Figure 8c).

One final consideration is related to the precision of innervation we report among secondary motor neurons. Much of our understanding of the molecular mechanisms of muscle targeting in zebrafish arises from work studying primary motor neurons (Beattie, 2000; Hutson and Chien, 2002; Lewis and Eisen, 2003), due to their discrete tiling of the axial musculature. We now show that there are secondary motor neurons with equally precise innervation patterns and others that are less specific but equally stereotyped. These differences in peripheral innervation are also mirrored by differences in central innervation, where motor neurons with less peripheral discrimination are also more likely to have recurrent collaterals. Since axon collaterals are concentrated in the neuropil and the dendrites of different units are intermingled, this suggests that a molecular recognition cues or activity levels rather than positioning is guiding the appropriate wiring of recurrent circuitry. Consequently, our findings not only provide a foundation for a future examination of the origins of recurrent motor circuitry, but also for work exploring the molecular and/or activity-dependent programs responsible for patterns of central and peripheral motor neuron innervation in both health and disease.

## Acknowledgements

We thank Elissa Szuter for fish care, Matthew Chiarelli for generating pilot data, and Eli Cadoff for help creating transgenic lines. We are also grateful to Dr. Juan Brusés for providing the mnx1:GFP construct, Dr. Andrew Miri for advice generating contour density plots, and members of the lab for helpful discussions and feedback. Financial support provided by The Graduate School, NIH R25 GM121231 and NIH R01 NS067299.

